# Investigations into the declining health of alder (*Alnus glutinosa*) along the river Lagan in Belfast, including the first report of *Phytophthora lacustris* causing disease of *Alnus* in Northern Ireland

**DOI:** 10.1101/2019.12.13.875229

**Authors:** Richard O Hanlon, Julia Wilson, Deborah Cox

## Abstract

Common alder (*Alnus glutinosa*) is an important tree species, especially in riparian and wet habitats. Alder is very common across Ireland and Northern Ireland, and provides a wide range of ecosystem services. Surveys along the river Lagan in Belfast, Northern Ireland led to the detection of several diseased *Alnus* trees. As it is known that Alnus suffers from a *Phytophthora* induced decline, this research set out to identify the presence and scale of the risk to *Alnus* health from *Phytophthora* and other closely related oomycetes. Sampling and a combination of morphological and molecular testing of symptomatic plant material and river baits identified the presence of several *Phytophthora* species, including *Phytophthora lacustris*. A survey of the tree vegetation along an 8.5 km stretch of the river revealed that of the 166 *Alnus* trees counted, 28 were severely defoliated/diseased and 9 were dead. Inoculation studies using potted *Alnus* saplings demonstrate that *P. lacustris* was able to cause disease, and Koch’s postulates for this pathogen-host combination were completed, which suggests a future risk to *Alnus* health from *P. lacustris* in Northern Ireland.

## Introduction

Common alder (*Alnus glutinosa* (L.) Gaertn.) is native to Europe, being common across Britain and Ireland (Clapham et al. 1952). Alder accounts for 2.7% of the forest estate in Ireland (NFI 2017), and is known as a relatively short lived tree (ca. 150 years; Mitchell 1996) that is suitable for planting in sites prone to water logging (Horgan et al. 2004). Native alder woods are common on wet poorly drained sites in Ireland, and can support a diverse herb and bryophyte layer (Cross 2012). Planting of *Alnus* in forests is supported in Ireland and Northern Ireland by government forestry grants (DAFM 2015; DAERA 2019). Furthermore, it has been suggested that introducing *Alnus* into plantations of Sitka spruce (*Picea sitchensis*) in Ireland and Britain would add structural diversity and have a positive effect on biodiversity and associated ecosystem services (Deal et al. 2014).

In 1993, a dieback of *Alnus* was noted in Britain, associated with a then unknown *Phytophthora* species (Gibbs et al. 1999). Further research identified the pathogen as being widespread in England and Wales (Streito 2003), and the pathogen was formally described as *Phytophthora alni* (Brasier et al. 2004). *Phytophthora alni sensu lato* was later split into three taxa, the species *Phytophthora uniformis*, and the hybrid species *Phytophthora* × *multiformis* and *Phytophthora* × *alni* based on molecular analysis (Husson et al. 2015). Dieback of alder caused by these pathogens has now been recorded in 17 European countries (Bjelke et al. 2016). Alder dieback was first confirmed in Ireland in 1999 (Clancy and Hamilton 1999), associated with *Phytophthora* × *multiformis* (O Hanlon et al. 2016a), while findings of alder dieback associated with *P. uniformis* were made in 2016 (O Hanlon et al. 2016b). To date, none of the previously mentioned pathogens have been detected in Northern Ireland (O Hanlon et al. 2016a), although dieback symptoms on alder have been evident for several years. For example, images on the free online resource Google street view from July 2008 to July 2018 show signs of alder dieback (e.g. thinning crown, bleeding cankers on the trunk, tree mortality) along the river Lagan in Belfast Northern Ireland UK (Figure 1). Haphazard surveys of the river bank of the Lagan close to the Agri-Food and Biosciences Institute headquarters in Newforge lane, Belfast have also identified *Alnus* trees with symptoms of Phytophthora infection (Fig. 2). Symptoms include thinning foliage composed of small yellowing leaves, and bleeding cankers on the trunk, similar to the symptoms normally caused by other tree infecting *Phytophthora* species (Jung et al. 2018).

**Figure 1.**
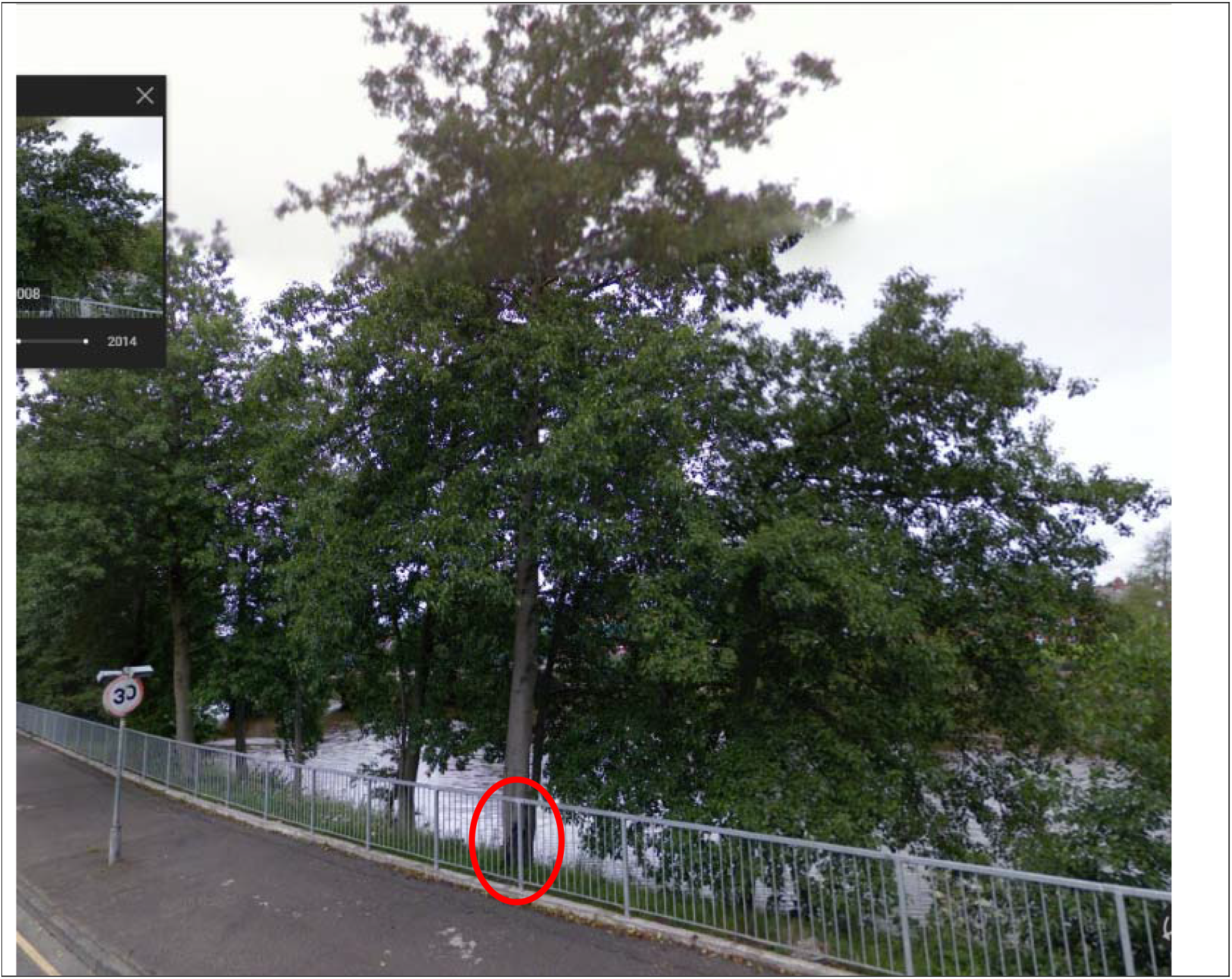
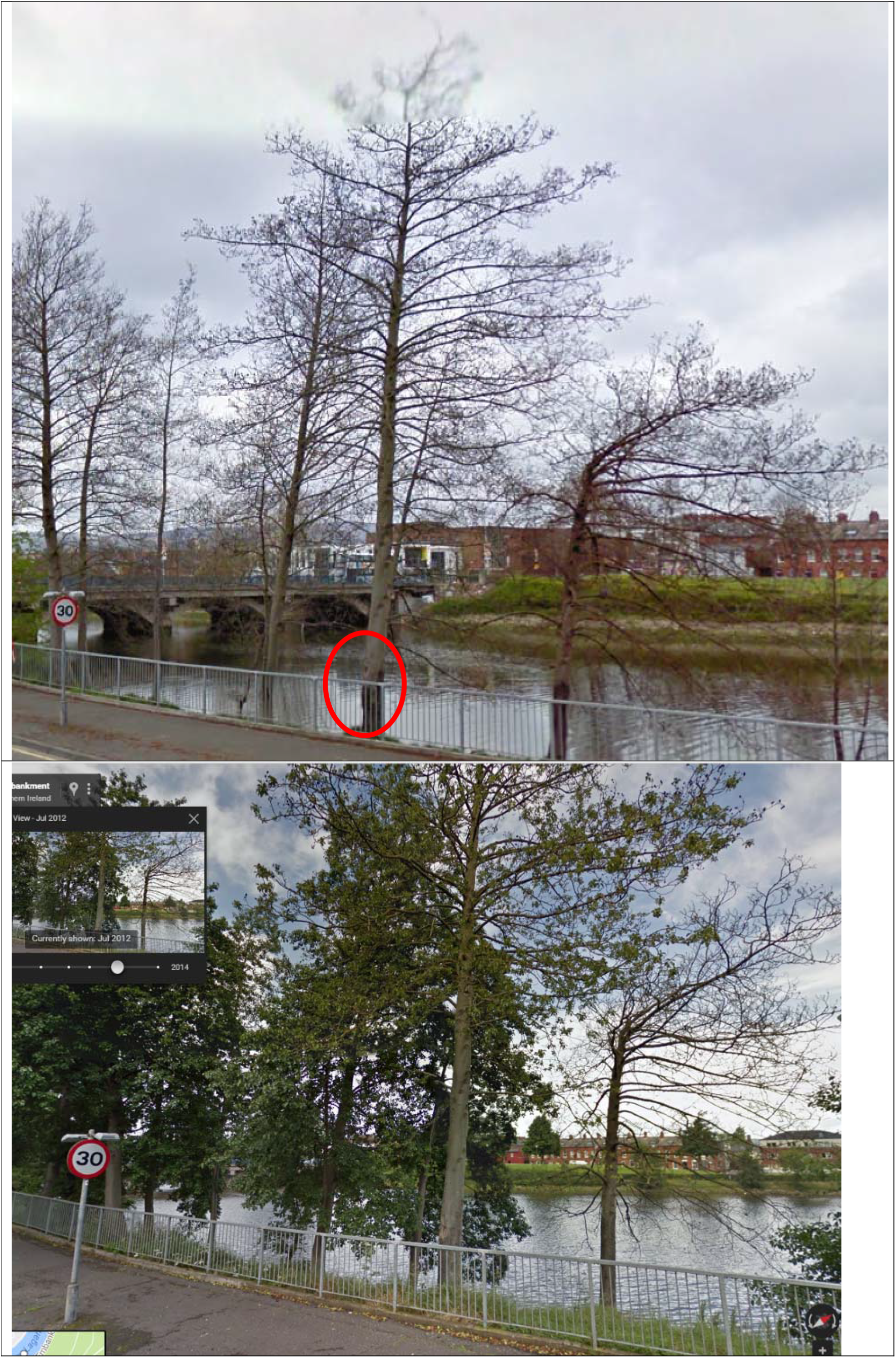
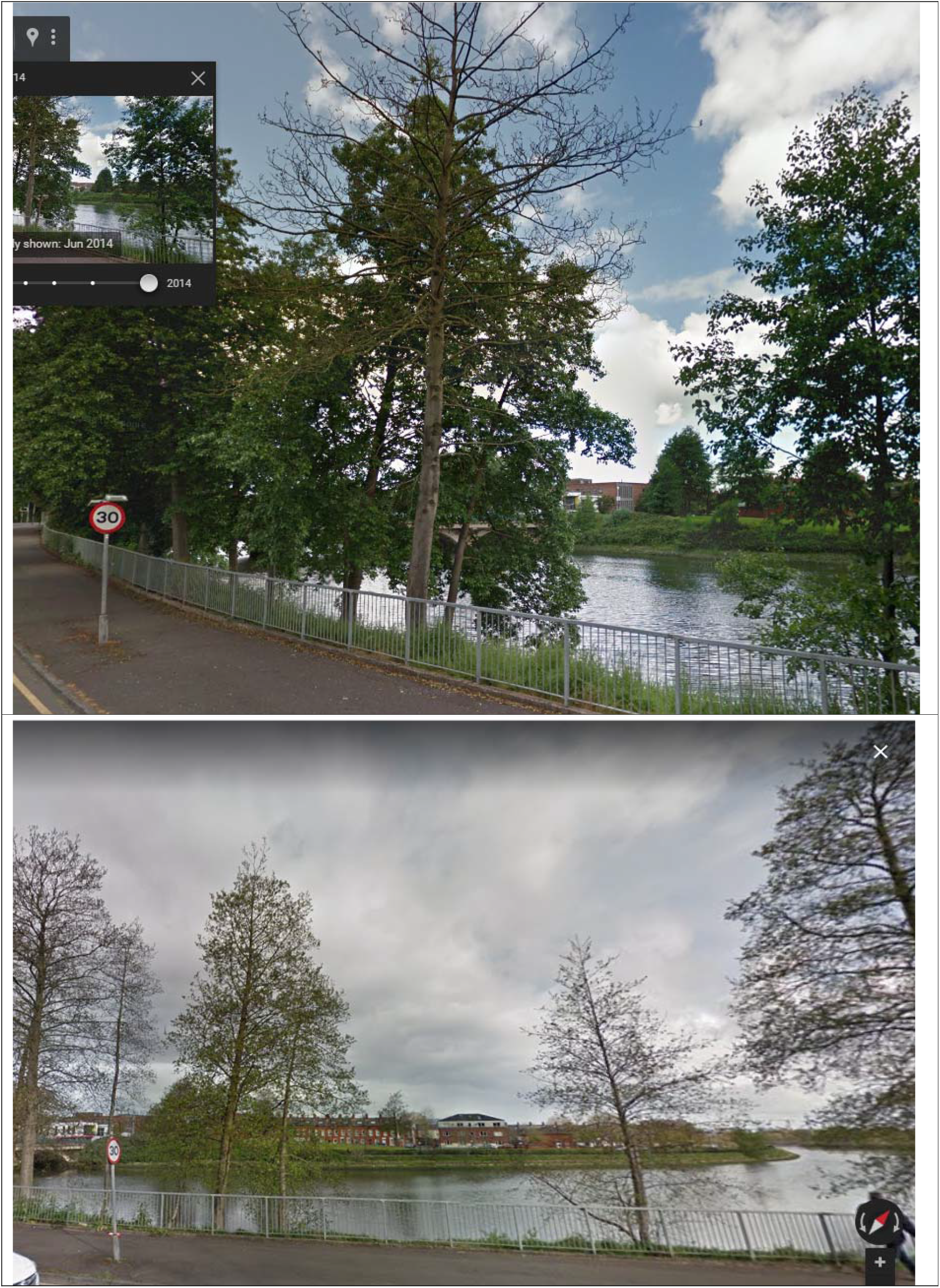
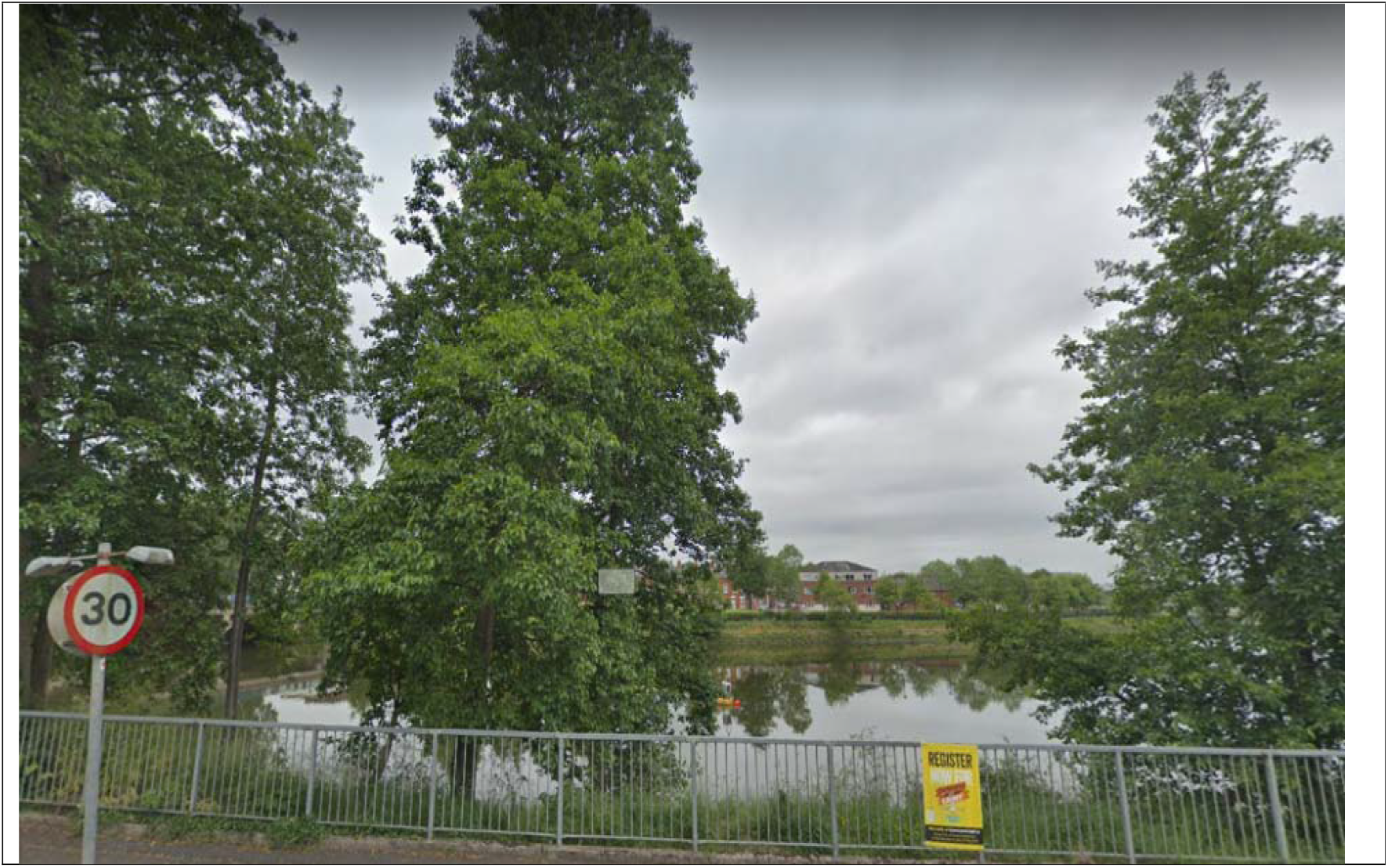
(composite) Chronosequence of Google street view images showing symptoms of alder dieback along the river Lagan in Belfast, Northern Ireland (54°34’58.1“N, 5°55’12.8”W). The images were taken in different months, starting from top to bottom: July 2008, April 2010, July 2012, June 2014, April 2017, and July 2018. Areas showing typical *Phytophthora* bleeding cankers have been encircled in a red oval.

**Figure 2:**
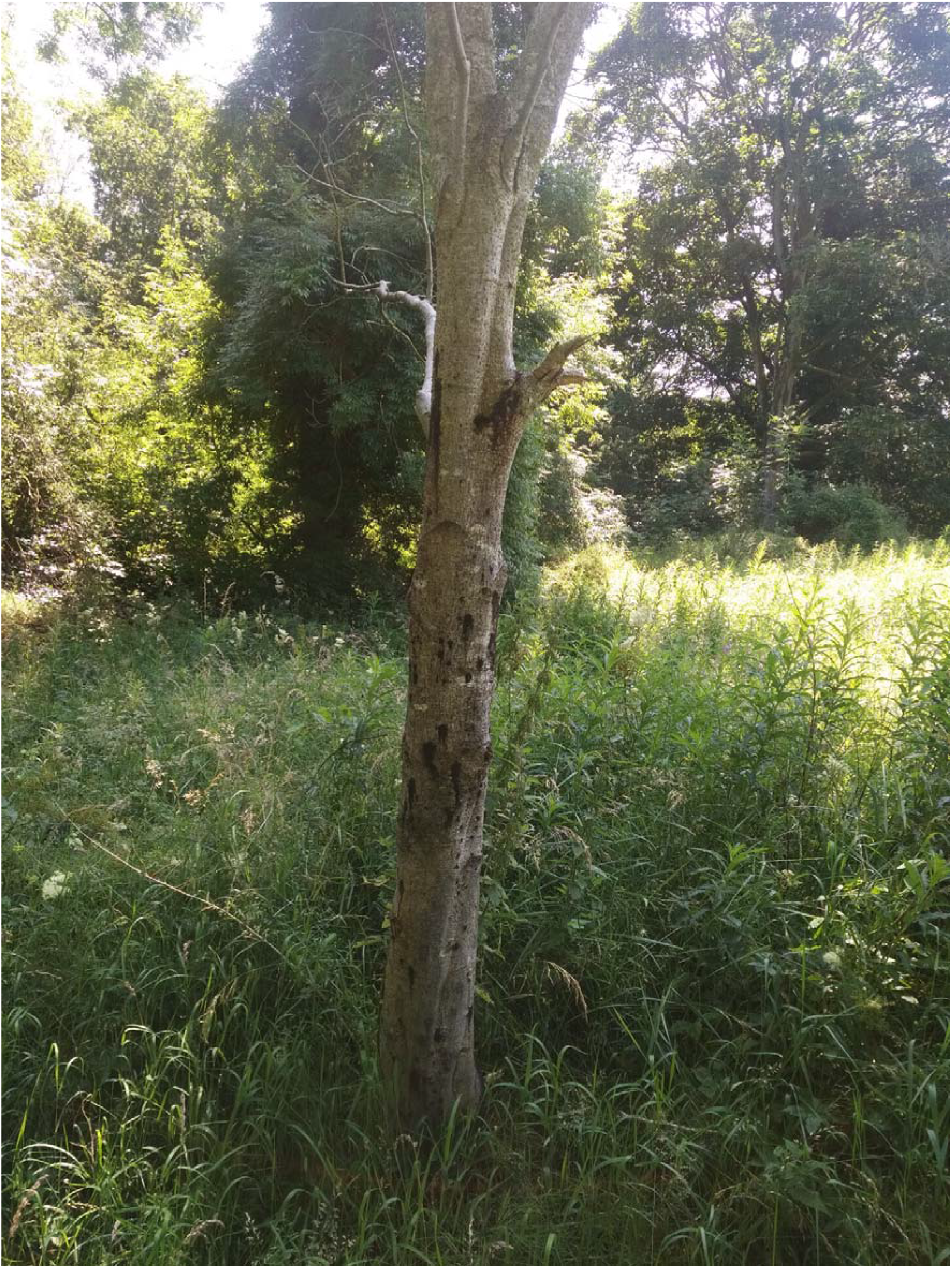
Bleeding cankers on *Alnus glutinosa* tree from which *Phytophthora lacustris* isolate P17-120 was isolated

The aim of this research was to investigate the health status of *Alnus* along the river Lagan, and to assess if *Phytophthora* species were impacting on *Alnus* health. This was assessed by (i) testing symptomatic trees for *Phytophthora* infection, (ii) conducting live plant inoculation studies to confirm the pathogenicity of *Phytophthora lacustris* to *Alnus glutinosa*, (iii) sampling the river using leaf baiting to identify if *Phytophthora* species were present in the water and (iv) carrying out an *Alnus* health survey along a portion of the river.

## Methods

### Symptomatic tree sampling and testing

In 2018, bark samples 1 cm think (including the cambium) from the upper and lower limits of bleeding cankers on a single dead *A. glutinosa* tree (Fig. 2; 54°33’26.9“N 5°56’10.7”W) were taken. Samples were surface sterilized and processed according to O Hanlon et al (2016b). Briefly, samples were put onto P5ARP[H] agar (1 litre distilled water, 17g cornmeal agar, 10 mg Pimiricin, 250 mg Ampicillin, 10 mg Rifamycin, 100 mg PCNB; Jeffers & Martin, 1986) and incubated at room temperature (19°C) for up to 2 weeks. After 3 days incubation at 19°C, *Phytophthora*–like mycelium was sub-cultured onto carrot piece agar (Werres et al. 2001) and incubated for another 5 days at 19°C. The isolate (P17-120) was stored in culture at 4°C for use in pathogenicity testing.

### Water baiting methods

At 2 points along the Lagan, leaf bait bags were deployed according to Turner et al., (2006) and O’Hanlon et al. (2018) with minor modification as follows: A leaf of *Alnus glutinosa*, *Rhododendron ponticum*, and *Quercus petraea* was removed from live, symptomless (i.e. putatively disease free) plants on the day of baiting, and sealed into a bait bag. The bait bag was composed of a double layer of muslin cloth, which was stapled shut, tied to the river bank with fishing line, and dropped into the river and left to float in the current. Bait bags were collected after 1 week, returned to the lab and the leaves removed from the bag. The leaves were processed in a similar way to the symptomatic tree samples (above). *Phytophthora*-like isolates produced from baits were grouped into morphogroups based on colony morphology and spore dimensions (Gallegly & Hong, 2008). When sporangia did not form on carrot piece agar, pieces of the outer colony edge were cut from the agar and put into a clean petri dish, and flooded with non-sterile pond water (i.e. a mixture of 50% water from the river Lagan and 50% tap water) to induce spore formation according to Scanu et al. (2015). The pond water was >6 months old at the time, in order to provide some assurance that any *Phytophthora* spores that may have been in the water were dead. Zappia et al. (2014) review oomycete risks from water and indicate that in general, survival for more than a few days is uncommon. A representative of each morphogroup was identified by DNA sequencing of the ITS gene region using the primers ITS4 and ITS5 primers (White et al. 1990). The DNA sequence was checked against the database of curated *Phytophthora* sequences in PhytophthoraID (Grünwald, et al. 2011), and also against the uncurated reference database available in GenBank using the Basic Local Alignment Search Tool (BLAST) (Altschul et al. 1990). The isolate from the symptomatic Alnus tree (P17-120) was deposited in GenBank under the accession number MH784622.

### Pathogenicity testing

Potted seedlings of alder (<3 years old) that were visually disease free were purchased from Coillte nurseries in Co Carlow, of which 12 were inoculated with agar plugs taken from the edge of the *P. lacustris* colony (P17-120) growing on carrot agar. Five other seedlings were inoculated with sterile carrot agar plugs to act as a negative control. In July 2018, each plant was wounded with a 3cm cut into the bark approximately halfway up the stem, with the agar plug inserted into the cut and sealed with damp cotton wool and parafilm. The plants were kept outside in a dedicated experimental area and watered regularly. After 6 weeks the outer bark was carefully scraped off and the lesions examined and measured. Data on lesion lengths and treatment was plotted and compared for differences between treatments (inoculation vs control) according to the method of Ho et al. (2019) using the online tool available at https://www.estimationstats.com. The entire diseased area in each plant was removed with a sterile knife, and transferred to the laboratory for *Phytophthora* isolation as per above methodology.

### Riparian tree visual survey

In order to investigate the extent of potential *Alnus* decline along the river, a visual survey of tree conditions was carried out along an 8.5 km stretch of the river. Mortality (dead, living), the level crown defoliation class (1= low defoliation, 4= extensive defoliation; Lakatos et al. 2014), age group (1= Diameter at Breast Height (DBH) < 10cm, 2= DBH 10 – 20, 3= DBH>20) and the presence of cankers for every *Alnus* tree was recorded. Notes were also taken of other tree health aspects for the surveyed *Alnus* trees. In order to get a rough estimation of the other tree species present in the area, the neighbouring species of tree to the left and right each *Alnus* tree was also recorded, if that tree was not also an *Alnus*.

## Results

### Symptomatic plant sampling

The isolate (P17-120) from the symptomatic *Alnus* tree was fast growing and produced a petaloid colony morphology similar to *Phytophthora gonapodyides*. The isolate produced non-caducous, non-papillate, mainly ovoid (some obpyriform) sporangia (average 35 × 25 μm). Sporangia showed internal and external proliferation and often had wide exit pores. These characteristics are in line with the description of *P. lacustris* (Nechwatal et al. 2013). Comparison of the DNA sequence from this isolate with PhytophthoraID indicated that the isolate was a close match (99%) to *Phytophthora lacustris* (Table 1).

**Table 1.** Results of the DNA sequencing of representative isolates of each of the morphogroups.

| **Sample** | **Best match**<br>**PhytophthoraID**<br>**(% similarity)** | **Best match**<br>**GenBank**<br>**(% similarity)** | **No. isolates**<br>**collected** |
| --- | --- | --- | --- |
| Symptomatic *Alnus*<br>canker (P17-120) | *Phytophthora lacustris*<br>(99) | *P. lacustris* (100) | 1 |
| Baiting morphotype 1 | *P. lacustris* (99) | *Phytophthora*<br>*gonapodyides* (96) /<br>*P. lacustris* (96) | 3 |
| Baiting morphotype 2 | *Phytophthora chlamydospora* (99) | *P. chlamydospora* (99) / *P. lacustris* (99) | 13 |
| Baiting morphotype 3 | *P. lacustris* (97) | *P. lacustris* (97) / *P. gonapodyides* (97) | 91 |
| Baiting morphotype 4 | *Pythium vexans* (80) | *Phytopythium litorale* (99) | 52 |

### Leaf baiting

Over 10 weeks of baiting, with weekly collections from 2 locations, 159 isolates were collected. These isolates were grouped into 4 morphogroups based on gross colony morphology, and spore microscopic characteristics. DNA sequencing indicated that these morphogroups represented 4 different taxa, although % similarity between the isolates collected and those in the online databases varied (Table 1). In three cases (Morphotype 1, 2, 3) the isolates were equally similar to two different species in the Genbank database. Morphotype 3 was commonly isolated, with morphotype 1 only isolated on 3 occasions.

### Pathogenicity testing

After 6 weeks dark lesions around the inoculation point were observed. The plants inoculated with *P. lacustris* isolate P17-120 had lesions of mean length 4.4 cm (n=12, SD+/− 0.76) whereas control plants had lesions of mean length 3.42 (n=5, SD+/− 0.60) (Fig. 3). This difference in lesion size related to treatment was confirmed by a Mann Whitney U test (U= 51.5, p<0.05). Attempts to isolate *Phytophthora* from the wounding points on inoculated and control saplings were carried out, with *P. lacustris* (confirmed by morphology) re-isolated from the necrotic tissue from all of the inoculated trees. No *Phytophthora* cultures were isolated from control plants. This completes Koch’s postulates for this host-pathogen pair in Northern Ireland.

**Figure 3.**
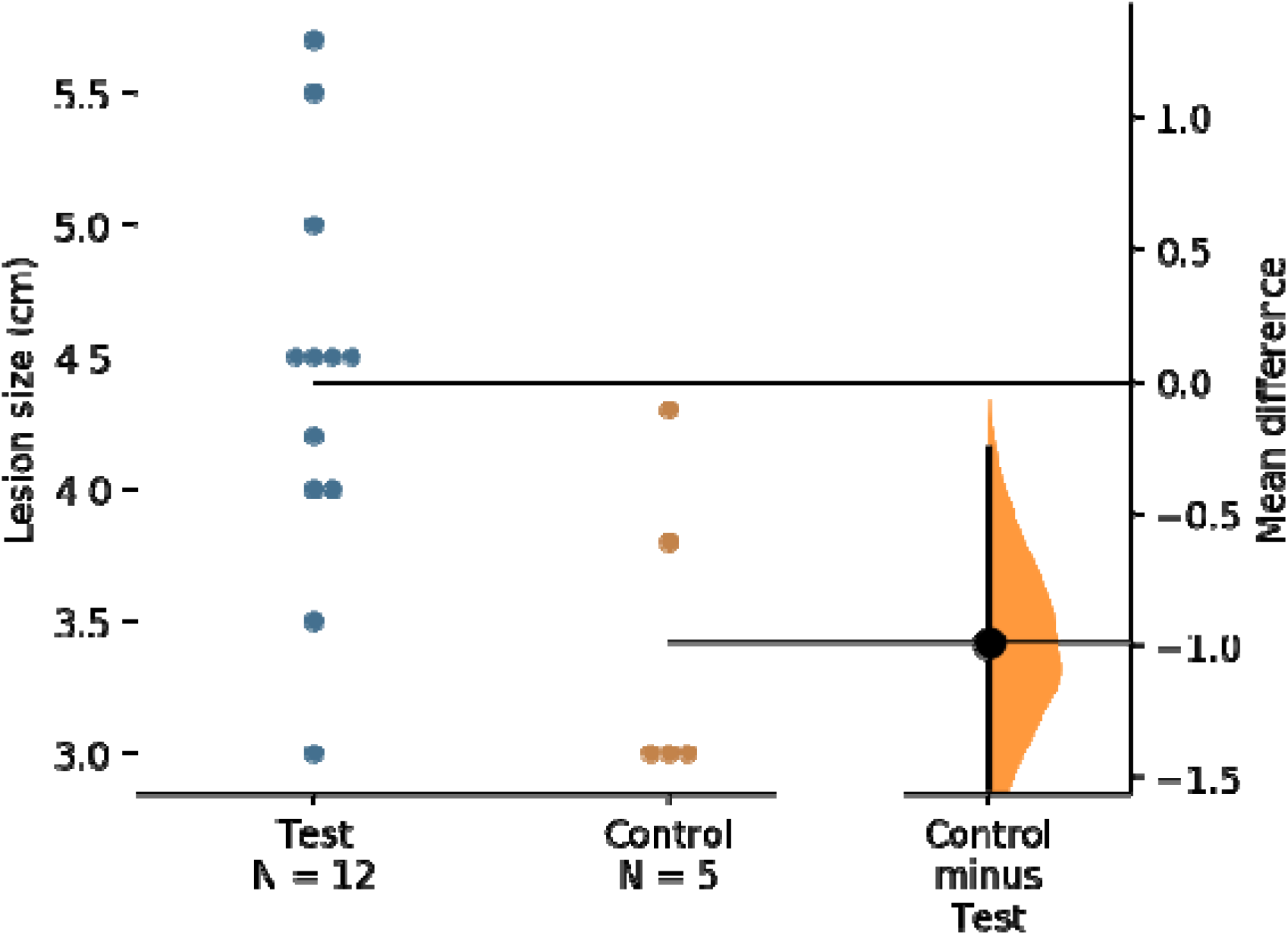
Estimation plots of data form the pathogenicity test. Saplings were inoculated with *Phytophthora lacustris* (test) or with sterile agar plugs (control). The filled curve indicates the complete difference in means distribution, given the observed data. The 95% confidence interval of difference in means is illustrated by the thick black line. The difference in means between the treatments (inoculation with *P. lacustris* or control) was significantly different based on a Mann Whitney U test (U= 51.5, p<0.05).

### Alder survey

A total of 166 *Alnus* trees were surveyed along an 8.5 km stretch on one side of the Lagan. The majority (83% or 137 trees) had diameters greater than 10 cm (Supplementary table 1). Of the total number of alder, 121 had low defoliation (1 or 2), with 28 showing moderate to severe defoliation. Nine *Alnus* trees were dead. Nine of the trees had bleeding cankers, and all of these were in defoliation class 3 or 4. Other tree species recorded were *Fraxinus excelsior* (n=23), *Salix* spp. (14), *Acer pseudoplatanus* (9), *Crataegus monogyna* (8), *Quercus* spp. (5), *Betula* spp. (3), *Fagus sylvatica* (3) and *Populus* spp. (3). There was one tree each of *Tilia cordata* and *Corylus avellana* recorded.

## Discussion

Alder dieback due to *Phytophthora* is currently widespread along river systems in many European countries including southern Germany (Jung and Blaschke 2004), Northern France (Thoirain et al. 2007), Czech Republic (Cerny and Strnadova 2010), Sweden (Redondo et al. 2015), and England (Webber et al. 2004). The disease is also present in several other countries in Europe (Jung et al. 2018), including Ireland (O’Hanlon et al. 2016a), though large scale surveys are needed to delimit its extent. The large scale death of *Alnus* in these river systems will likely have many negative impacts on the ecosystems services these trees provide (Cerny et al. 2008; Bjelke et al. 2016). Alder has many important functions in riparian habitats, including provision of shade, Nitrogen fixation, stabilization of river banks through its root network, provision of habitat for many organisms, and increasing the aesthetic appeal of river banks (Bjelke et al. 2016). In Ireland and Northern Ireland, it is the 6^th^ most commonly recorded tree species in the wild, after *Crataegus* spp., *Fraxinus excelsior*, *Acer pseudoplatanus*, *Ilex aquifolium*, and *Prunus spinosa* (National Biodiversity Data Centre 2019).

Jung et al. (2018) provide a comprehensive review of the distribution of *Alnus* decline in Europe. They highlight that although the main cause of the decline is usually either *Phytophthora uniformis*, the hybrid taxa *Phytophthora* × *alni* or *Phytophthora* × *multiformis*; other *Phytophthora* species are also known to cause disease in *Alnus*. These include *Phytophthora cactorum*, *P. gonapodyides*, *P. lacustris*, *Phytophthora plurivora* and *Phytophthora polonica*. This research has found that *P. lacustris* is possibly responsible for *Alnus* decline along the river Lagan in Belfast, Northern Ireland. *Phytophthora lacustris* has also been found infecting *A. glutinosa* in Portugal (Kanoun-Boule et al. 2016), as well as several other plant hosts in Europe (Nechwatal et al. 2013). The river baiting carried out in the study also indicates that there is a risk of *P. lacustris* spreading along the river and infecting more trees. This is similar to findings of *P. lacustris* in soil and streams in Ireland (O’Hanlon et al. 2016b). It is well known that *Phytophthora* species spread readily in water in commercial (Zappia et al. 2014) and wider environment (Sims et al. 2014) locations. Gibbs et al. (1999) also found evidence that *Phytophthora* disease of *Alnus* in England was being spread in the river. Worryingly, the low number of young *Alnus* trees recorded in the survey may indicate that *P. lacustris* is reducing recruitment of young *Alnus* trees, as well as killing older *Alnus*. Younger plants are known to be more susceptible to *Phytophthora* infection than older ones (Oßwald et al. 2014). This could lead to a situation where *Alnus* becomes less common along the river Lagan, thus reducing the ecosystem services provided by *Alnus*.

This study used a combination of morphotyping and DNA sequencing to identify the oomycete diversity. Unfortunately, the ITS primers used did not allow for resolution of the species to a high degree (>99%) of confidence. There were also issues with the analysis returning two different species names of equal similarity to the morphotypes collected. O’Hanlon et al. (2016b) also discussed this issue, when their sequencing was unable to separate closely related taxa within *Phytophthora* clade 7. The use of several markers is encouraged to separate closely related species within the genus *Phytophthora* (Hyde et al., 2014; Yang and Hong 2018). The result of the DNA analyses also depended on the database used. With PhytophthoraID being a curated database including only sequences of deposited species confirmed by competent taxonomists (Grünwald, et al. 2011), this resource is likely to be more reliable and accurate. GenBank on the other hand is not curated, and is known to suffer from erroneous names of sequence deposits of other microorganisms (e.g. fungi; Nilsson, et al. 2006) in the database. It is highly likely that the four morphotypes identified from the baiting samples are *P. gonapodyides*, *P. chlamydospore*, *P. lacustris* and *Phytopythium litorale*, as suggested by the DNA sequencing. These species have been found in surveys in Ireland (O’Hanlon et al. 2016b), and also in other samples from watercourses in Northern Ireland (O’Hanlon 2017). They are also common in other regions of the world (Cerny et al. 2015; Duarte et al. 2015; Hansen et al. 2015). None of the species, except *P. lacustris*, is known as a threatening plant pathogen, and they probably function in speeding up the decay of plant material in streams (Hansen et al. 2012).

It is interesting to speculate if, and how, *P. lacustris* may have entered the area. It is not known if *P. lacustris* is native to Ireland and Northern Ireland, with Jung et al. (2016) unable to assign its putative native range. The species has been linked with plant trade infrequently across Europe (Jung et al. 2016) and the USA (Bienapfl and Balci 2014), and therefore could have been introduced inadvertently into Northern Ireland on infected plant material or soil. The area of the Lagan surveyed in this research is highly modified by humans, with the nearby Belvoir forest area having a long history of history of exotic plant introductions. The riverbank also has a widespread infestation of *Rhododendron ponticum*, Japanese knotweed (*Fallopia japonica*), Giant Hogweed (*Heracleum mantegazzianum*), and Himalayan balsam (*Impatiens glandulifera*). The nearby Belvoir forest also has had recent infestations of other important *Phytophthora* species, including the regulated *Phytophthora ramorum* on Japanese larch (*Larix kaempferi*) and the previously regulated *Phytophthora lateralis* on Lawson Cypress (*Chamaecyparis lawsoniana*). Coupled with this, there has been an invasion and outbreak of the non-native insect pests horsechestnut leaf miner (*Cameraria ohridella*) on horsechestnut (*Aesculus hippocastanum*) (Anon 2014) and ash sawfly *Tomostethus nigritus* on ash *F. excelsior* in the area since 2016 (Jess et al. 2017).

This investigation is the first to demonstrate *P. lacustris* as a cause of disease in *Alnus* on the island of Ireland (i.e. in Northern Ireland or Ireland). Though we did not detect either of the other serious *Phytophthora* pathogens of *Alnus*, *P. uniformis* or *P.* × *alni*, it is also likely that they are responsible for *Alnus* decline in Northern Ireland, as these have been recorded in *Alnus* in Ireland (O’Hanlon et al. 2016a). The ecology of the river ecosystem of the river Lagan is under threat from *Phytophthora* and many other invasive species. Better biosecurity education will help prevent further introductions for invasive species to habitats in Ireland and Northern Ireland. There is also good potential to use regular river baiting and riparian tree surveys (or amateur records) as an early warning system for new plant pest invasions.

## Acknowledgements

The authors acknowledge the Royal Society of Biology, DEFRA, BSPP and N8 AgriFood for funding JW on a Plant Health Studentship. The research was funded by the Department of Agriculture, Environment and Rural Affairs Northern Ireland.

---

**Supplementary table 1.** *Alnus* tree health survey results. The level crown defoliation class (1= low defoliation, 4= extensive defoliation; Lakatos et al. 2014), age group (1= Diameter at Breast Height (DBH) < 10cm, 2= DBH 10 – 20, 3= DBH >20) and the presence of cankers for every *Alnus* tree was recorded. Notes were also taken of other tree health aspects for the surveyed *Alnus* trees.

| Alder identifier | Defoliation class score | Age | Presence of cankers | notes |
| --- | --- | --- | --- | --- |
| A1 | 4 | 2 | 0 |  |
| A2 | 0 | 2 | 0 |  |
| A3 | 1 | 2 | 0 |  |
| A4 | 3 | 2 | 1 |  |
| A5 | 3 | 2 | 1 |  |
| A6 | 0 | 2 | 0 |  |
| A7 | 0 | 1 | 0 |  |
| A8 | 0 | 2 | 0 |  |
| A9 | 0 | 2 | 0 |  |
| A10 | 4 | 1 | 1 |  |
| A11 | 0 | 2 | 0 |  |
| A12 | 0 | 1 | 0 |  |
| A13 | 2 | 2 | 0 |  |
| A14 | 0 | 2 | 0 |  |
| A15 | 0 | 2 | 0 |  |
| A16 | 0 | 3 | 0 |  |
| A17 | 0 | 2 | 0 |  |
| A18 | 0 | 2 | 0 |  |
| A19 | 0 | 2 | 0 |  |
| A20 | 0 | 2 | 0 |  |
| A21 | 0 | 3 | 0 |  |
| A22 | 0 | 2 | 0 |  |
| A23 | 3 | 2 | 0 |  |
| A24 | 2 | 2 | 0 |  |
| A25 | 0 | 2 | 0 |  |
| A26 | 0 | 2 | 0 |  |
| A27 | 0 | 2 | 0 |  |
| A28 | 0 | 2 | 0 |  |
| A29 | 0 | 2 | 0 |  |
| A30 | 2 | 2 | 0 | Galls |
| A31 | 0 | 2 | 0 |  |
| A32 | 0 | 2 | 0 |  |
| A33 | 0 | 2 | 0 |  |
| A34 | 0 | 2 | 0 |  |
| A35 | 0 | 2 | 0 |  |
| A36 | 0 | 2 | 0 |  |
| A37 | 0 | 2 | 0 |  |
| A38 | 3 | 2 | 0 | almost dead |
| A39 | 0 | 3 | 0 |  |
| A40 | 4 | 3 | 0 | lots of bracket fungi |
| A41 | 0 | 2 | 0 |  |
| A42 | 0 | 2 | 0 |  |
| A43 | 0 | 2 | 0 |  |
| A44 | 0 | 2 | 0 |  |
| A45 | 0 | 2 | 0 |  |
| A46 | 0 | 2 | 0 |  |
| A47 | 0 | 2 | 0 |  |
| A48 | 0 | 2 | 0 |  |
| A49 | 0 | 2 | 0 |  |
| A50 | 3 | 2 | 0 | dead |
| A51 | 3 | 2 | 0 | dead |
| A52 | 3 | 2 | 0 | dead |
| A53 | 4 | 2 | 0 |  |
| A54 | 4 | 2 | 0 | ivy present |
| A55 | 0 | 2 | 0 |  |
| A56 | 0 | 2 | 0 |  |
| A57 | 0 | 2 | 0 |  |
| A58 | 0 | 2 | 0 |  |
| A59 | 0 | 2 | 0 |  |
| A60 | 0 | 2 | 0 |  |
| A61 | 0 | 3 | 0 |  |
| A62 | 0 | 3 | 0 |  |
| A63 | 3 | 3 | 0 | Dead on one side |
| A64 | 0 | 1 | 0 |  |
| A65 | 0 | 1 | 0 |  |
| A66 | 3 | 3 | 1 | dead crown. Girdled stem,<br>galls |
| A67 | 0 | 1 | 0 |  |
| A68 | 1 | 1 | 0 |  |
| A69 | 3 | 3 | 1 | mostly in the river |
| A70 | 3 | 1 | 0 | almost dead |
| A71 | 0 | 1 | 0 |  |
| A72 | 0 | 1 | 0 |  |
| A73 | 1 | 3 | 0 | Ivy present |
| A74 | 0 | 1 | 0 |  |
| A75 | 0 | 1 | 0 |  |
| A76 | 0 | 1 | 0 |  |
| A77 | 0 | 1 | 0 |  |
| A78 | 0 | 1 | 0 |  |
| A79 | 0 | 1 | 0 |  |
| A80 | 0 | 1 | 0 |  |
| A81 | 1 | 1 | 0 |  |
| A82 | 0 | 2 | 0 |  |
| A83 | 0 | 3 | 0 |  |
| A84 | 0 | 2 | 0 |  |
| A85 | 3 | 1 | 1 |  |
| A86 | 1 | 3 | 0 |  |
| A87 | 1 | 1 | 0 |  |
| A88 | 0 | 1 | 0 |  |
| A89 | 0 | 2 | 0 |  |
| A90 | 2 | 1 | 0 |  |
| A91 | 0 | 2 | 0 |  |
| A92 | 2 | 1 | 0 |  |
| A93 | 1 | 2 | 0 |  |
| A94 | 2 | 2 | 0 |  |
| A95 | 0 | 2 | 0 |  |
| A96 | 0 | 2 | 0 |  |
| A97 | 0 | 2 | 0 |  |
| A98 | 2 | 2 | 0 |  |
| A99 | 0 | 3 | 0 |  |
| A100 | 0 | 2 | 0 |  |
| A101 | 0 | 2 | 0 |  |
| A102 | 0 | 2 | 0 |  |
| A103 | 0 | 2 | 0 |  |
| A104 | 0 | 2 | 0 |  |
| A105 | 0 | 2 | 0 |  |
| A106 | 0 | 2 | 0 |  |
| A107 | 0 | 2 | 0 |  |
| A108 | 0 | 2 | 0 |  |
| A109 | 0 | 2 | 0 |  |
| A110 | 0 | 2 | 0 |  |
| A111 | 0 | 2 | 0 |  |
| A112 | 0 | 2 | 0 |  |
| A113 | 0 | 2 | 0 |  |
| A114 | 0 | 2 | 0 |  |
| A115 | 0 | 2 | 0 |  |
| A116 | 0 | 2 | 0 |  |
| A117 | 0 | 2 | 0 |  |
| A118 | 0 | 2 | 0 |  |
| A119 | 0 | 1 | 0 |  |
| A120 | 0 | 2 | 0 |  |
| A121 | 0 | 3 | 0 |  |
| A122 | 0 | 3 | 0 |  |
| A123 | 2 | 2 | 0 | Ivy present, tree 40% dead |
| A124 | 1 | 2 | 0 |  |
| A125 | 0 | 2 | 0 |  |
| A126 | 0 | 2 | 0 |  |
| A127 | 0 | 2 | 0 |  |
| A128 | 2 | 2 | 0 |  |
| A129 | 0 | 2 | 0 |  |
| A130 | 0 | 2 | 0 |  |
| A131 | 0 | 2 | 0 |  |
| A132 | 0 | 2 | 0 |  |
| A133 | 0 | 2 | 0 |  |
| A134 | 0 | 2 | 0 |  |
| A135 | 0 | 2 | 0 |  |
| A136 | 0 | 2 | 0 |  |
| A137 | 0 | 2 | 0 |  |
| A138 | 0 | 2 | 0 |  |
| A139 | 0 | 2 | 0 |  |
| A140 | 0 | 2 | 0 |  |
| A141 | 0 | 2 | 0 |  |
| A142 | 0 | 2 | 0 |  |
| A143 | 0 | 2 | 0 |  |
| A144 | 0 | 2 | 0 |  |
| A145 | 2 | 2 | 0 |  |
| A146 | 3 | 2 | 0 |  |
| A147 | 0 | 2 | 0 |  |
| A148 | 0 | 2 | 0 |  |
| A149 | 0 | 2 | 0 |  |
| A150 | 2 | 2 | 0 |  |
| A151 | 0 | 1 | 0 |  |
| A152 | 4 | 2 | 0 |  |
| A153 | 2 | 2 | 0 |  |
| A154 | 2 | 2 | 0 |  |
| A155 | 0 | 1 | 0 |  |
| A156 | 0 | 1 | 0 |  |
| A157 | 3 | 2 | 0 | almost dead |
| A158 | 0 | 2 | 0 |  |
| A159 | 0 | 2 | 0 |  |
| A160 | 0 | 2 | 0 |  |
| A161 | 2 | 3 | 0 |  |
| A162 | 0 | 1 | 0 |  |
| A163 | 0 | 1 | 0 |  |
| A164 | 4 | 2 | 1 |  |
| A165 | 4 | 2 | 1 |  |
| A166 | 4 | 2 | 1 |  |

